# Manganese Ion Inhibits Influenza A Virus Replication by Targeting the PA Endonuclease

**DOI:** 10.64898/2026.07.27.740892

**Authors:** Yueqi Wu, Shan Xu, Yigang Tong

## Abstract

Influenza A virus remains a major global health threat, with rapid mutation and emerging drug resistance underscoring the urgent need for novel antivirals. Manganese ions (Mn^2+^) are known to enhance host antiviral immunity, but their direct effects on influenza virus replication remain elusive. Here, we demonstrate that manganese chloride (MnCl_2_) potently inhibits the replication of both H1N1 and H3N2 subtypes in cultured cells at micromolar concentrations, with antiviral activity not shared by other divalent cations (Ca^2+^, Cu^2+^, Zn^2+^, Mg^2+^). Mechanistically, MnCl_2_ acts predominantly at post-entry stages, suppresses viral mRNA synthesis, and directly inhibits the RNA cleavage activity of the PA endonuclease. Pharmacological antagonism with the PA inhibitor baloxavir further supports PA as a key target. In a mouse model, intranasal MnCl_2_ administration alleviated body weight loss, reduced mortality, and decreased pulmonary viral RNA loads. Collectively, these findings identify Mn^2+^ as a direct inhibitor of influenza virus replication that acts, at least in part, by targeting the PA endonuclease, providing a conceptual framework for metal ion-based antiviral strategies.

## Introduction

Influenza virus remains one of the most persistent viral pathogens, causing annual seasonal epidemics and occasional pandemics. In a typical year, seasonal influenza is estimated to cause up to 5 million severe cases and 500,000 deaths globally (Fineberg, 2014). The 2009 H1N1 pandemic underscored the vulnerability of public health systems and the urgent need for novel antivirals. Clinically approved antiviral agents for influenza primarily include M2 ion channel blockers (e.g., amantadine), neuraminidase inhibitors (e.g., oseltamivir, zanamivir), and the polymerase inhibitor baloxavir marboxil(Ngiam *et al*, 2026; Pomeshikova *et al*, 2026; Sato, 2025). However, the widespread use of these agents has led to significant drug resistance, including M2 blocker resistance in nearly all circulating influenza A virus strains(Sato, 2025), oseltamivir resistance (H275Y in N1 neuraminidase), and baloxavir resistance (I38T in PA), highlighting the urgent need for additional antiviral strategies(Omoto *et al*, 2018; Collins *et al*, 2009).

Among the proteins encoded by influenza virus, the RNA-dependent RNA polymerase (RdRp) represents a highly promising antiviral target owing to its central role in viral transcription and replication and its high degree of conservation across subtypes (Honda & Ishihama, 1997; te Velthuis & Fodor, 2016). The RdRp consists of three subunits: PA, PB1, and PB2. The PB1 subunit harbors the polymerase catalytic center responsible for RNA chain elongation; the PB2 subunit binds the cap structure of host pre-mRNAs to initiate cap-snatching; and the N-terminal domain of the PA subunit (PAN) cleaves host pre-mRNAs to generate capped primers for viral transcription(Lamb & Choppin, 1976; Sugiyama *et al*, 2009). Crystal structures of PAN solved by Dias et al. and Yuan et al. in 2009 revealed that the endonuclease active site contains conserved divalent metal ion-coordinating motifs and is highly conserved across influenza subtypes (Dias *et al*, 2009; Yuan *et al*, 2009). These features have established PA endonuclease as an ideal target for anti-influenza drug development, a notion strongly validated by the successful approval of baloxavir marboxil as the first PA endonuclease-targeting anti-influenza drug(Li *et al*, 2026; DuBois *et al*, 2012).

Metal ions play critical roles in virus–host interactions(Charlie-Silva *et al*, 2019). Manganese (Mn), an indispensable trace element, is essential for metabolic regulation, enzymatic catalysis, and immune homeostasis(Fijałkowski *et al*, 2025; Coradduzza *et al*, 2025). While its role as an essential nutrient has been well established in the context of host physiology(Qin & Xu, 2026), a growing body of evidence has revealed its direct antiviral functions at the molecular level. Mn^2+^ enhances cGAS sensitivity to dsDNA, promoting cGAS–STING activation and host defense against DNA viruses(Wang *et al*, 2018). Beyond this pathway, Mn^2+^ exerts broad-spectrum antiviral activity against RNA viruses (PRRSV, VSV) and DNA viruses (HSV-1) in a cGAS-STING-independent manner(Sun *et al*, 2023). Mn^2+^ also enhances type I interferon production (IFN-α, IFN-β, IFN-λ1) and promotes antiviral immunity via TBK1 phosphorylation(Sui *et al*, 2022, 2025). Furthermore, Mn^2+^ directly suppresses viruses such as FMDV and HBV, and modulates lysosomal and oxidative stress responses during infection(Zhang *et al*, 2023). Collectively, these findings establish Mn^2+^ as both an immune modulator and a direct antiviral agent.

Notably, the active site of influenza virus PA endonuclease is precisely a catalytic pocket that depends on the coordination of divalent metal ions(Doan *et al*, 1999). Biochemical studies have shown that PA endonuclease exhibits substantially higher affinity for Mn^2+^ than for Mg^2+^, and that different divalent metal ions can induce distinct conformational states that affect catalytic efficiency and substrate preference (Noble *et al*, 2012; Crépin *et al*, 2010). These findings suggest that the availability and local concentration of Mn^2+^ may directly influence PA endonuclease function and, consequently, viral polymerase activity. However, whether and how Mn^2+^ regulates influenza virus replication through direct modulation of PA endonuclease activity remains largely unexplored.

In this study, we combined cellular, biochemical, and in vivo approaches to investigate the inhibitory effect of Mn^2+^ on H1N1 replication, focusing on its direct impact on PA endonuclease activity. Our findings provide new mechanistic insights into metal ion–polymerase interactions and establish a conceptual foundation for metal-based antiviral strategies.

## Results

### MnCl_2_ inhibits influenza A virus replication in cell culture

To evaluate the antiviral activity of manganese chloride (MnCl_2_) against influenza A virus, we first assessed its cytotoxicity and antiviral efficacy in MDCK and A549 cells. Cell viability assays showed that MnCl_2_ exhibited minimal cytotoxicity within the concentration range used in this study. The half-maximal cytotoxic concentration (CC_50_) of Mn^2+^ exceeded 200 μM in both MDCK and A549 cells **(Fig. 1, Fig. S1)**, indicating a favorable safety margin for subsequent antiviral analyses.

**Figure 1.**
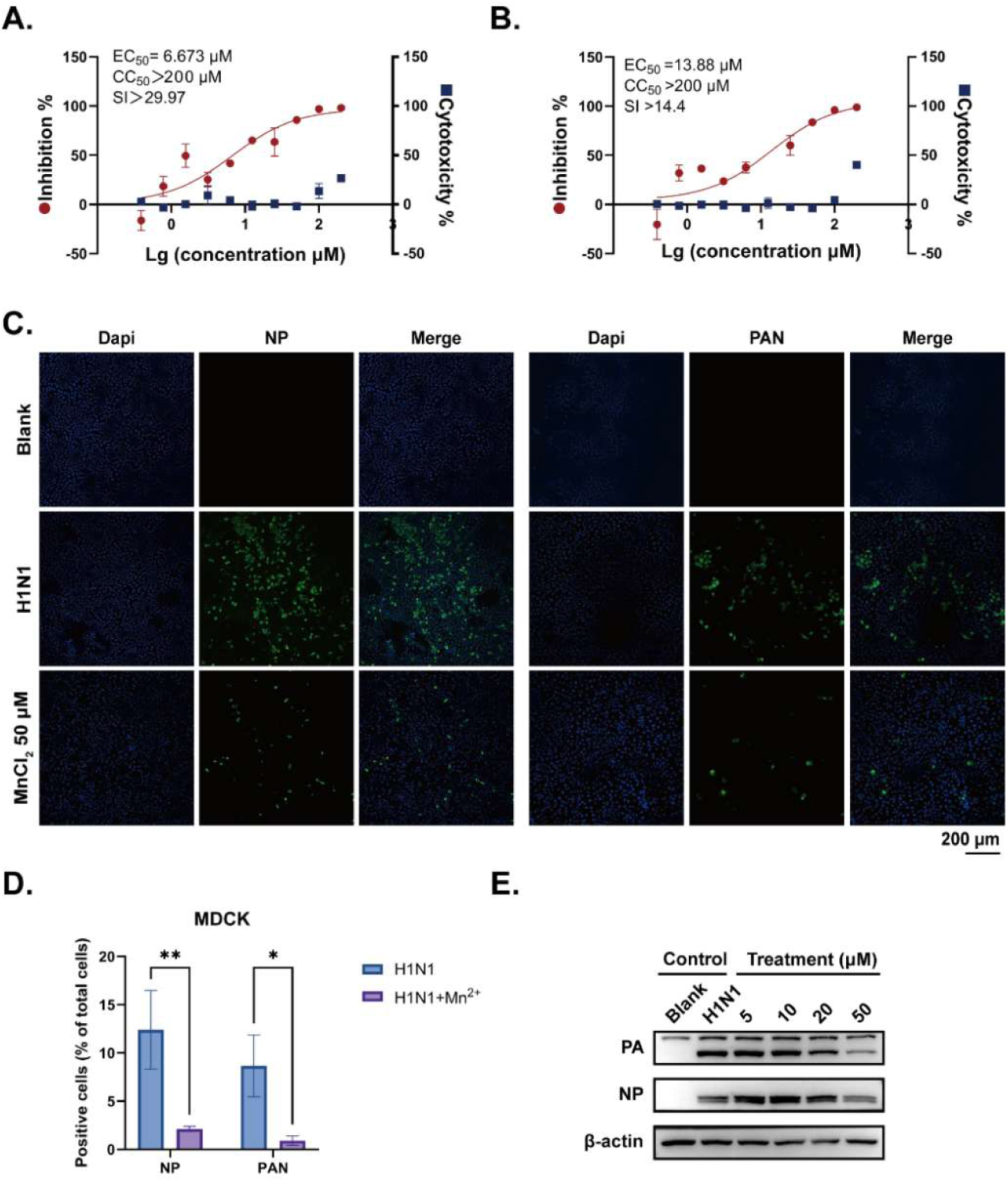
Mn^2+^ inhibits H1N1 and H3N2 replication in MDCK cells. (A) Antiviral activity (red) and cytotoxicity (blue) of Mn^2+^ against H1N1 in MDCK cells. EC_50_, CC_50_, and SI values are indicated. (B) Antiviral activity (red) and cytotoxicity (blue) of Mn^2+^ against H3N2 in MDCK cells. (C) Immunofluorescence staining of viral NP (green) and PA (green) in H1N1-infected MDCK cells with or without Mn^2+^ (50 μM) treatment. Nuclei were counterstained with DAPI (blue). Scale bar = 200 μm. (D) Quantification of NP- and PA-positive cells from (C). (E) Western blot analysis of NP and PA protein levels in H1N1-infected MDCK cells treated with Mn^2+^ (5-50 μM). β-actin served as loading control. **Data information:** In (A, B, D), data are presented as mean ± SD (n = 3). ****P < 0.0001 versus virus-infected control (Student’s t-test).

We next examined the antiviral efficacy of MnCl_2_ against both H1N1 and H3N2 influenza A viruses in MDCK cells. MnCl_2_ treatment resulted in a significant and dose-dependent reduction in the replication of both viral subtypes. In MDCK cells, the EC_50_ of MnCl_2_ was 6.67 μM for H1N1 and 13.88 μM for H3N2 **(Fig. 1A, B)**. The antiviral activity was further confirmed in A549 cells, where MnCl_2_ also suppressed H1N1 and H3N2 replication in a dose-dependent manner, with EC_50_ values not exceeding 100 μM **(Fig. S1)**. In contrast, other divalent metal ions (Ca^2+^, Cu^2+^, Zn^2+^, and Mg^2+^) tested under identical conditions showed no appreciable anti-influenza activity in MDCK cells **(Fig. S2)**. These results demonstrate that MnCl_2_ broadly inhibits influenza A virus replication across different viral subtypes and in distinct cellular contexts, with a degree of metal ion selectivity.

To further validate the inhibitory effect of MnCl_2_ at the protein level, viral protein expression was examined by immunofluorescence and Western blot analyses using antibodies against NP and PA. In MDCK cells, MnCl_2_ treatment markedly reduced the proportion of both NP- and PA-positive cells **(Fig. 1C, D)**, and Western blot analysis confirmed a substantial decrease in intracellular NP and PA levels **(Fig. 1E)**. Consistent results were obtained in A549 cells **(Fig. S1C–E)**, further supporting the antiviral efficacy of MnCl_2_ across different cell types.

### MnCl_2_ acts at a post-entry stage of the influenza virus life cycle

To determine the stage of the influenza virus life cycle targeted by MnCl_2_, a time-of-addition (TOA) assay was performed with MnCl_2_ added at distinct phases of infection, as illustrated in **Fig. 2A**. When MnCl_2_ was pre-incubated with H1N1 virions prior to infection and removed after viral adsorption, no significant reduction in viral replication was observed compared with the virus control, indicating that MnCl_2_ does not directly inactivate viral particles. Similarly, MnCl_2_ treatment restricted to the viral adsorption and entry phase had minimal impact on subsequent viral replication.

**Figure 2.**
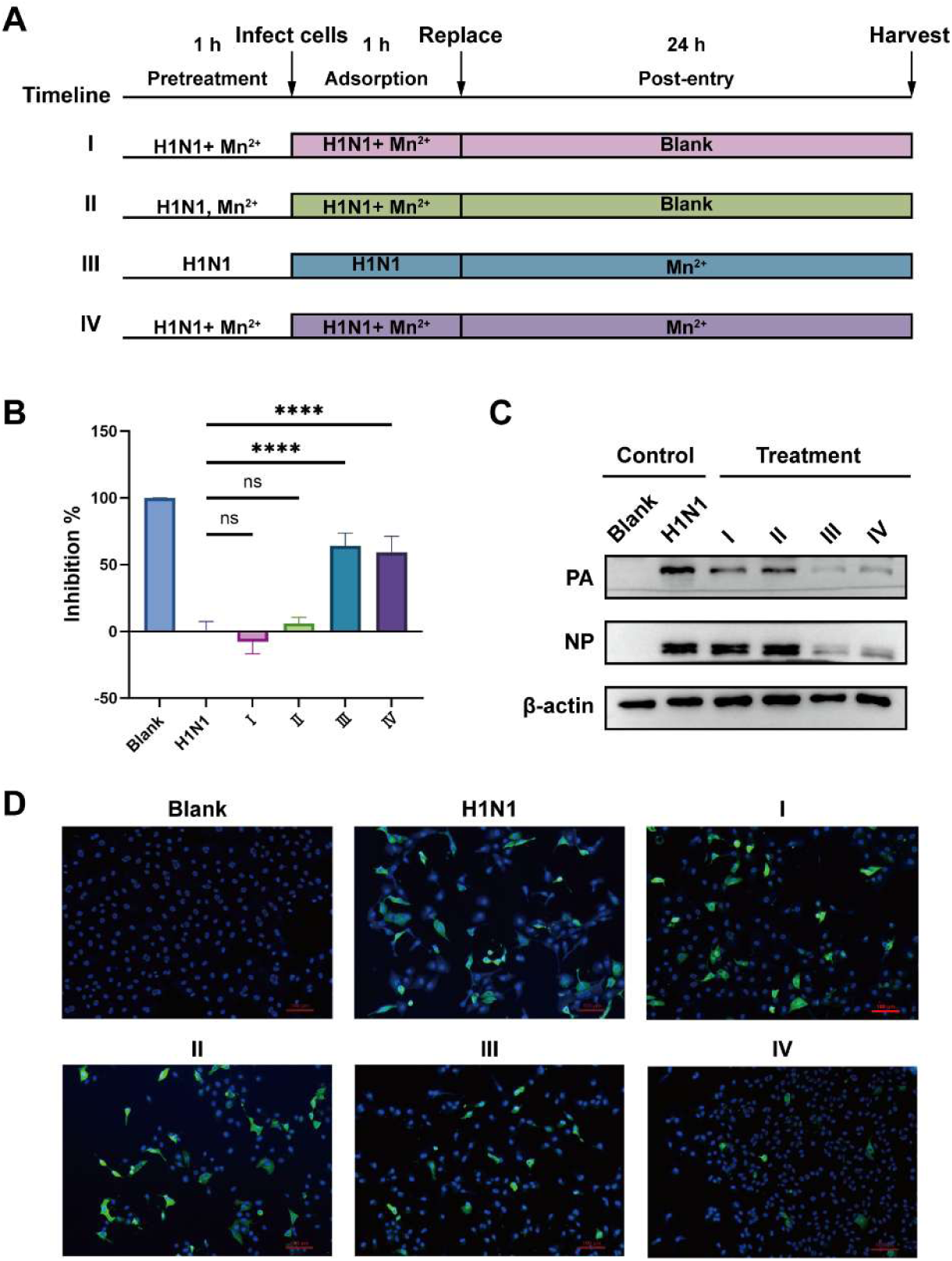
**Mn**^2+^ **acts at the post-entry stage of H1N1 replication. (A)** Schematic of the time-of-addition experimental design. MDCK cells were exposed to Mn^2+^ (50 μM) and H1N1 (100 TCID_50_) at different time windows as indicated. **(B)** RT-qPCR analysis of intracellular H1N1 RNA levels at 24 h post-infection. **(C)** Western blot analysis of viral NP and PA protein levels in the different treatment groups. β-actin served as the loading control. **(D)** Immunofluorescence staining of viral NP protein (green) in the different treatment groups. Nuclei were counterstained with DAPI (blue). Scale bar = 100 μm. **Data information:** In (B), data are presented as mean ± SD (n = 3). ****P < 0.0001 versus the virus-infected control; n.s., not significant (Student’s t-test).

In contrast, addition of MnCl_2_ after viral entry resulted in a marked suppression of virus replication. RT-qPCR analysis revealed that post-entry treatment significantly reduced viral RNA levels (****P < 0.0001 versus virus-infected control), whereas pretreatment or treatment limited to the adsorption phase had no significant effect (n.s.; **Fig. 2B**). Consistently, Western blot analysis showed that viral NP and PA protein levels were markedly decreased in the post-entry and full-time treatment groups **(Fig. 2C)**. Immunofluorescence staining further confirmed a visible reduction in the number of NP-positive cells under these conditions compared with the virus-infected control **(Fig. 2D)**.

Collectively, these results demonstrate that MnCl_2_ does not exert its antiviral activity by directly neutralizing influenza virions or blocking viral entry, but instead acts predominantly at a post-entry stage of infection, consistent with an effect on intracellular viral replication processes.

### MnCl_2_ suppresses early viral RNA accumulation following infection

To further dissect the stage at which MnCl_2_ interferes with the viral life cycle, we quantified the kinetics of viral mRNA, cRNA, and vRNA in the supernatant of MDCK cells at 2, 4, and 12 hpi, with baloxavir marboxil (BXM) as a polymerase inhibitor control **(Fig. 3A)**. MnCl_2_, like BXM, elicited sustained suppression of both mRNA and cRNA at all time points (****P < 0.0001 versus virus control). In contrast, vRNA reduction was transient, significant only at 4 hpi and largely lost by 12 hpi. This kinetic pattern argues against a block in virus assembly or release, which would produce sustained vRNA suppression at late time points, and instead indicates that MnCl_2_ restricts the cRNA template supply, thereby reducing the vRNA pool available for packaging while the release process itself remains intact.

**Figure 3.**
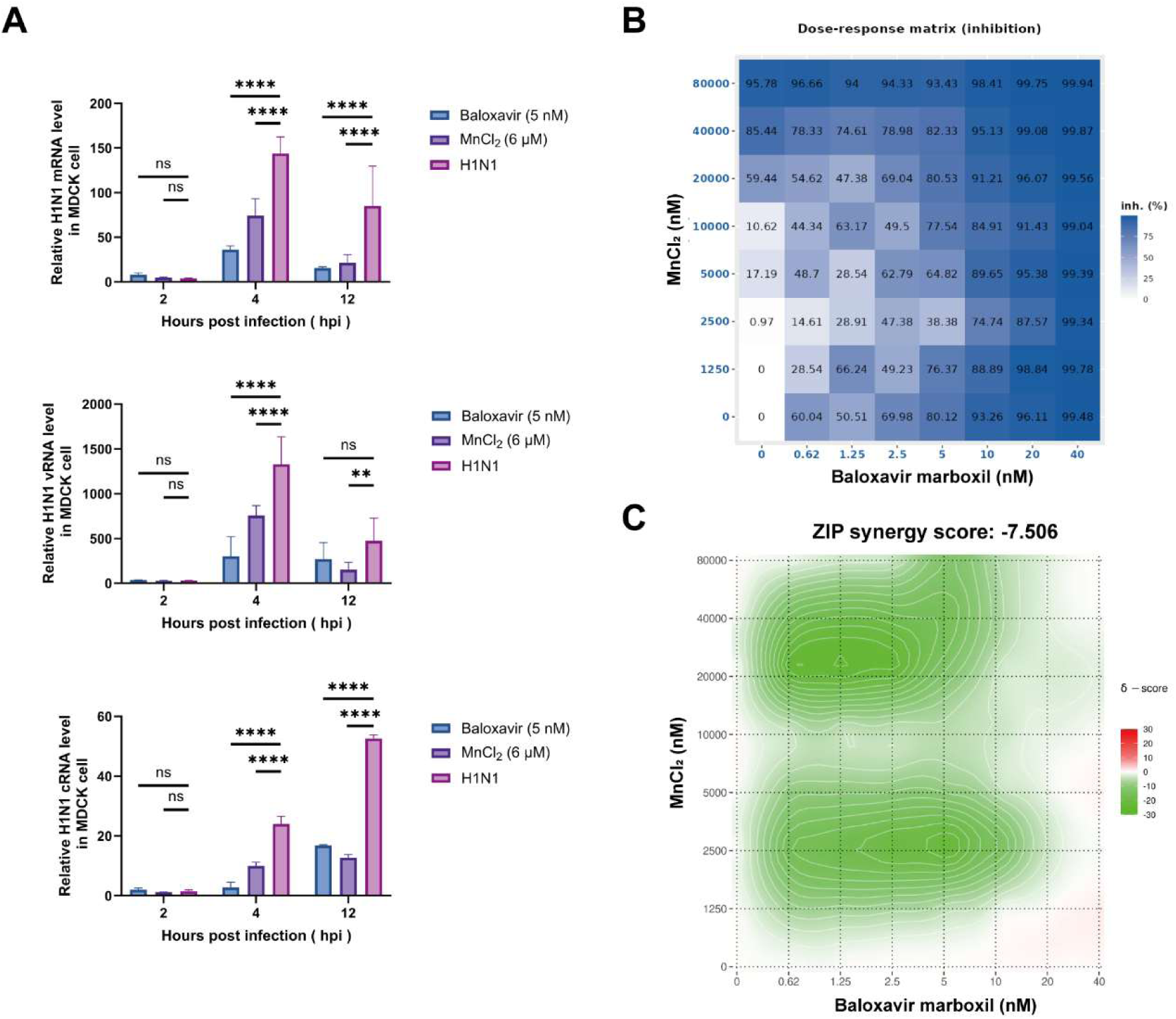
**Mn**^2+^ **suppresses H1N1 viral RNA synthesis and exhibits target antagonism with the PA inhibitor baloxavir.** (A) Kinetics of intracellular H1N1 mRNA, vRNA, and cRNA levels following Mn^2+^ treatment, determined by strand-specific RT-qPCR. (B) Heatmap of inhibition rates from an 8 × 8 checkerboard combination of Mn^2+^ and baloxavir against H1N1. Color intensity represents the percentage of viral inhibition. (C) ZIP synergy score heatmap derived from the data in (B). The x- and y-axes represent the concentrations of Mn^2+^ and baloxavir, respectively. The δ value indicates the deviation from the expected additive effect: δ > 0, synergy; δ ≈ 0, additivity; δ < 0, antagonism. Pronounced antagonism (δ < 0) was observed across the major pharmacologically active concentration ranges. **Data information:** In (A), data are presented as mean ± SD (n = 3).

The sustained suppression of mRNA and cRNA pinpoints the target of MnCl_2_ to early viral RNA synthesis. Persistent blockade of cRNA, the obligate template for vRNA amplification, demonstrates that MnCl_2_ impairs the catalytic activity of the viral replication complex. The close phenotypic concordance with BXM further supports a polymerase inhibitor-like mechanism.

To test this, we performed MnCl_2_–BXM combination experiments and evaluated their interaction by ZIP synergy analysis. Across an 8 × 8 checkerboard matrix, ZIP analysis revealed no synergy; instead, pronounced antagonism (δ < 0) was observed across the major active concentration ranges **(Fig. 3B, C)**, suggesting that both compounds target the same or overlapping sites, most likely the PA subunit of the viral RdRp complex. This pharmacological antagonism independently corroborates the RNA kinetics data.

Together with the time-of-addition results, these data demonstrate that MnCl_2_ targets the early post-entry transcription–replication axis of influenza virus by inhibiting mRNA transcription and cRNA replication, without interfering with budding or release. The kinetic dissociation between sustained mRNA and cRNA suppression and transient vRNA reduction, combined with the pharmacological antagonism with BXM, indicates that MnCl_2_ acts at the level of viral RNA polymerase activity, most likely by engaging the PA endonuclease active site.

### MnCl_2_ inhibits the RNA cleavage activity of the influenza virus PA endonuclease

To determine whether MnCl_2_ directly affects the enzymatic activity of the influenza virus PA endonuclease, recombinant PA protein was expressed and purified to near homogeneity, as confirmed by Coomassie brilliant blue staining **(Fig. 4A)**, and its identity was verified by Western blot analysis using an anti-PA antibody **(Fig. 4B)**. The purified PA protein was then subjected to in vitro RNA cleavage assays.

**Figure 4.**
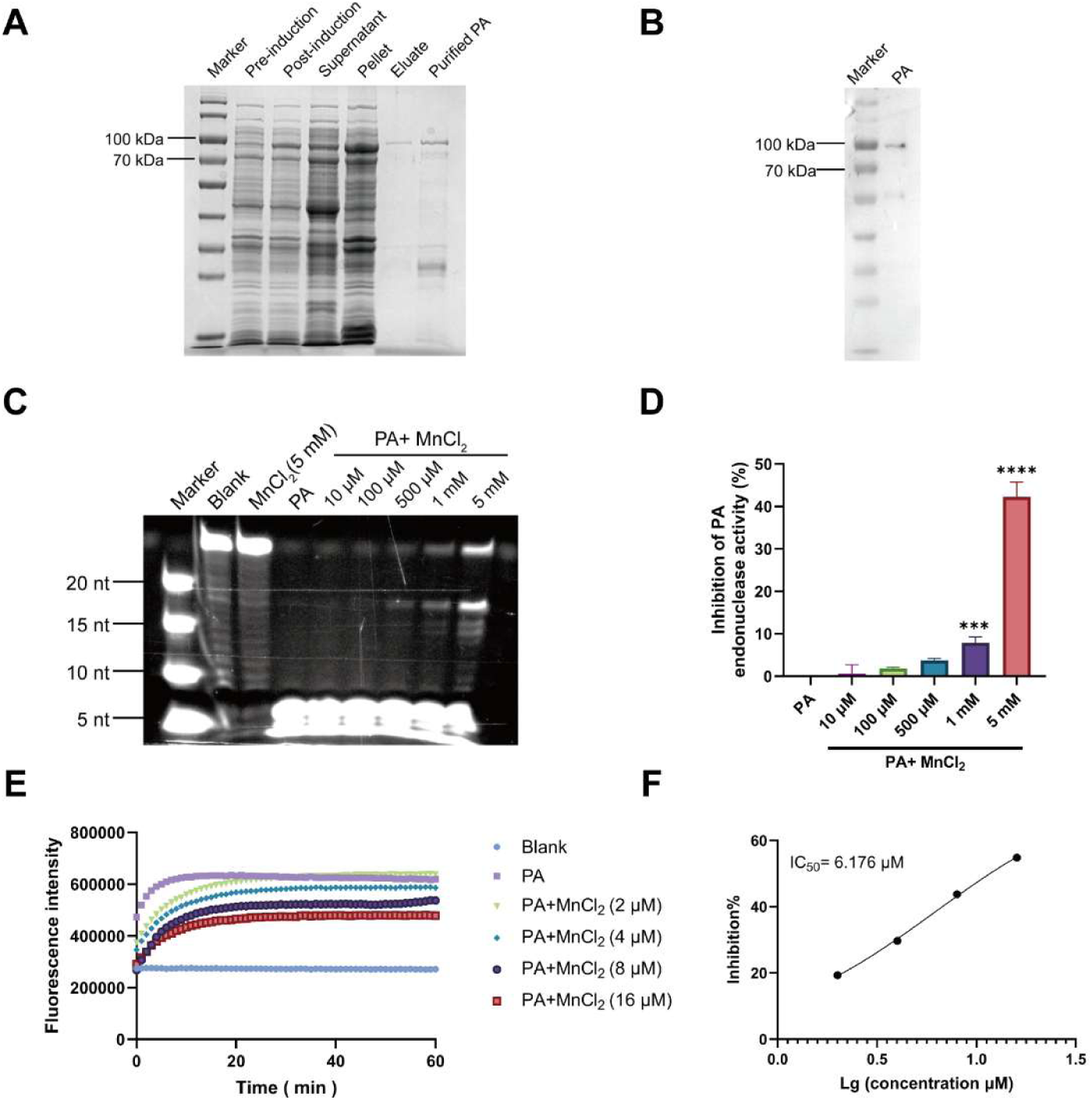
**Mn**^2+^ **directly inhibits the endonuclease activity of H1N1 PA protein in vitro.** (A) Coomassie brilliant blue staining of fractions from different purification stages and the final purified PA protein. (B) Western blot validation of the purified PA protein using an anti-PA antibody. The sample was diluted 20-fold before loading, with the same protein input as used in the RNA gel assay and fluorescence-based enzyme assay. (C) RNA gel analysis of PA endonuclease inhibition by Mn^2+^. Purified PA protein was pre-incubated with the indicated concentrations of MnCl_2_, followed by addition of fluorophore-labeled RNA substrate. Substrate-only control and MnCl_2_-only control (5 mM) were included as negative controls. (D) Inhibition rates of PA endonuclease activity at the indicated MnCl_2_ concentrations. (E) Real-time fluorescence-based monitoring of PA endonuclease activity. Fluorescence intensity was continuously recorded after substrate addition. Curves represent the cleavage kinetics at different MnCl_2_ concentrations. (F) Inhibition curve of Mn^2+^ against PA endonuclease activity. Reaction rates were calculated from the fluorescence signals at the 10-min time point in (E). IC_50_ was determined by nonlinear regression. **Data information:** In (D–F), data are presented as mean ± SD (n = 3). In (D), group comparisons were performed by one-way ANOVA. ***P < 0.001, ****P < 0.0001 versus PA-only control.

We first employed a fluorescence-based endonuclease activity assay using a dual-labeled RNA substrate (5′-FAM / 3′-BHQ1), in which fluorescence is quenched in the intact substrate and released upon PA-mediated cleavage. PA protein was pre-incubated with increasing concentrations of MnCl_2_, and the fluorescence signal was monitored in real time. MnCl_2_ treatment resulted in a concentration-dependent suppression of fluorescence increase, indicating impaired PA-mediated substrate cleavage **(Fig. 4E)**. The inhibition rate was calculated at the 10-minute time point as follows: Inhibition (%) = (Mean_sample_ − Mean_pa_) / (Mean_blank_ − Mean_pa_) × 100%, where Mean_pa_ represents the untreated PA control (0% inhibition) and Mean_blank_ represents the substrate-only control (100% inhibition). MnCl_2_ inhibited PA endonuclease activity in a dose-dependent manner, yielding an IC_50_ of 6.176 μM **(Fig. 4F)**.

To corroborate these findings with a gel-based readout, we performed an RNA cleavage assay in which the reaction products were resolved by denaturing PAGE. Consistent with the fluorescence-based results, MnCl_2_ markedly reduced the generation of cleaved RNA products in a concentration-dependent manner **(Fig. 4C)**. Quantification of the cleavage product band intensities confirmed significant inhibition at 1 mM MnCl_2_ (*P < 0.001) and further enhancement at 5 mM (**P < 0.0001), as determined by one-way ANOVA (Fig. 4D). Notably, although statistically significant inhibition was observed in the RNA gel assay at 1 mM and 5 mM MnCl_2_ **(Fig. 4D)**, the inhibition rates did not reach 50%, likely reflecting the lower sensitivity of the gel-based endpoint readout compared with the real-time fluorescence assay **(Fig. 4E, F)**.

To further verify the direct regulatory role of Mn^2+^ on PA endonuclease activity, we performed an EDTA chelation–rescue assay. PA protein was pre-treated with EDTA to chelate metal ions from its active site, which led to a marked decrease in enzymatic activity, as reflected by substantially reduced fluorescence signals. The subsequent addition of 100 μM or 500 μM MnCl_2_ fully restored PA endonuclease activity, with fluorescence curves rising to levels comparable to those of the untreated PA control (**Fig. S3**). In contrast, the addition of 1 mM MnCl_2_ failed to rescue enzymatic activity, and the fluorescence signal remained at a low level. These results demonstrate that Mn^2+^ at appropriate concentrations serves as an essential cofactor for PA endonuclease activity, whereas excess Mn^2+^ inhibits its catalytic function, consistent with the concentration-dependent inhibition of PA by Mn^2+^ described above.

Together, these results demonstrate that MnCl_2_ directly inhibits the RNA cleavage activity of the influenza virus PA endonuclease in vitro, supporting a role for Mn^2+^ in modulating viral polymerase function.

### MnCl_2_ reduces influenza virus–induced morbidity and viral burden in mice

To evaluate the antiviral efficacy of MnCl_2_ in vivo, mice were infected with a lethal dose of H1N1 and treated with MnCl_2_ or baloxavir at the time of infection. Compared with the virus-infected control group, Mn^2+^-treated mice exhibited markedly attenuated body weight loss **(Fig. 5A)** and significantly improved survival **(Fig. 5B)**. The baloxavir-treated group displayed the strongest protective effect, while the Mn^2+^-treated group, although slightly less potent, still conferred significant survival benefit.

**Figure 5.**
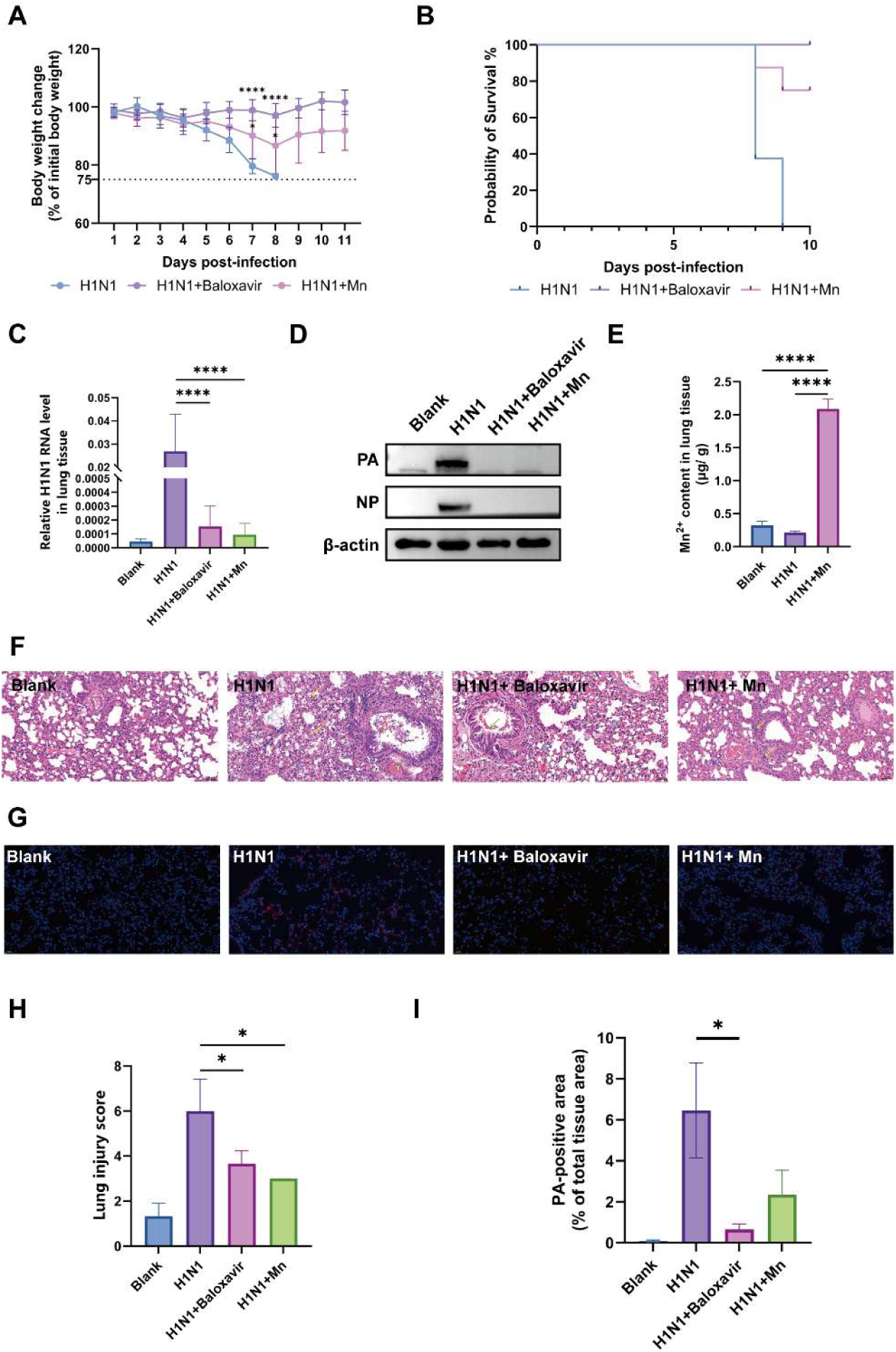
**Mn**^2+^ **suppresses H1N1 replication and attenuates lung injury in mice.** (A) Body weight changes over 11 days post-infection, normalized to the initial body weight on day 0. * indicates maximal divergence between curves. (B) Survival curves over 10 days post-infection. (C) Relative H1N1 RNA levels in lung tissues at day 5 post-infection, determined by RT-qPCR. (D) Western blot analysis of NP and PA protein levels in lung tissues at day 5 post-infection. β-actin served as loading control. (E) Mn^2+^ content in lung tissues at day 2 post-infection, measured by ICP-MS. (F) Representative H&E-stained lung sections at day 5 post-infection. Scale bar = 200 μm. Blue arrows: granulocyte infiltration in alveolar walls; yellow arrows: intra-alveolar lymphocyte and macrophage exudation; cyan arrows: eosinophilic exudate in alveolar spaces; orange arrows: necrotic debris in bronchiolar lumina; green arrows: vascular congestion. (G) Representative immunofluorescence staining of PA protein (green) in lung tissues. Nuclei were counterstained with DAPI (blue). Scale bar = 200 μm. (H) Semi-quantitative lung injury scores, calculated as the sum of scores for alveolar wall thickening, inflammatory cell infiltration, and necrosis. (I) Quantification of PA-positive area relative to total tissue area from (G). **Data information:** In (A, C, E, H, I), data are presented as mean ± SD (n = 3–5 mice per group). In (B), group comparisons were performed using the Log-rank (Mantel-Cox) test. In (C, E, H, I), **P < 0.01, ***P < 0.001 versus virus-infected control (one-way ANOVA with Tukey’s post-hoc test).

To assess the effect of MnCl_2_ on viral replication in the lung, viral RNA and protein levels were examined at day 5 post-infection. Mn^2+^ treatment significantly reduced pulmonary viral RNA levels **(Fig. 5C)** and markedly suppressed viral NP and PA protein expression **(Fig. 5D)**, indicating that Mn^2+^ effectively inhibits H1N1 gene expression and possesses clear antiviral activity. Consistent with these findings, intranasal administration of MnCl_2_ led to a significant enrichment of Mn in the lung tissue at day 2 post-infection, as determined by ICP-MS **(Fig. 5E)**, demonstrating efficient delivery of Mn^2+^ to the target organ.

Histopathological examination of lung sections revealed that virus-infected mice developed severe alveolar damage with extensive inflammatory infiltration, whereas Mn^2+^ treatment substantially attenuated these pathological changes **(Fig. 5F)**. Semi-quantitative lung injury scoring confirmed significantly reduced injury scores in the Mn^2+^-treated group **(Fig. 5H)**. Immunofluorescence staining further demonstrated a marked reduction of viral PA protein signals in the lungs of Mn^2+^-treated mice **(Fig. 5G)**. Quantification of the PA-positive area showed a trend toward reduction in the Mn^2+^-treated group compared with the virus-infected control, although the difference did not reach statistical significance **(Fig. 5I)**.

Collectively, these results demonstrate that intranasal MnCl_2_ administration effectively suppresses H1N1 replication in vivo, attenuates virus-induced lung pathology, and improves survival outcome in mice.

## Discussion

This study identifies Mn^2+^ as a direct inhibitor of influenza A virus replication and provides a mechanistic framework linking Mn^2+^ exposure to viral polymerase dysfunction. Through integrated cellular, biochemical, and in vivo analyses, we demonstrate that MnCl_2_ exerts potent antiviral activity against both H1N1 and H3N2 subtypes in MDCK and A549 cells, thereby expanding the functional repertoire of metal ions in influenza virus biology. The consistent antiviral efficacy observed across distinct viral subtypes and cell types suggests a broad-spectrum inhibitory mechanism that is neither subtype-specific nor cell-type-restricted.

Importantly, this antiviral activity was not a general property of divalent metal ions. Preliminary screening of selected divalent cations (Ca^2+^, Cu^2+^, Zn^2+^, and Mg^2+^) under identical conditions revealed no appreciable inhibition of H1N1 replication in MDCK cells **(Fig. S2)**, indicating a degree of metal ion selectivity.

Our time-of-addition experiments demonstrate that Mn^2+^ does not directly inactivate virions but acts at a post-entry stage, consistent with the pronounced suppression of early viral RNA accumulation. Given the essential role of PA-mediated cap-snatching in viral transcription and the high sequence conservation of the PA active site across influenza A subtypes, these results point to the viral polymerase as the likely target of Mn^2+^ interference. Consistent with this interpretation, ZIP analysis revealed antagonistic interactions between MnCl_2_ and the PA inhibitor baloxavir, independently supporting overlapping functional targets within the viral polymerase.

Biochemical assays using purified PA protein directly support this hypothesis. MnCl_2_ inhibited the RNA endonuclease activity of PA in a concentration-dependent manner, identifying the PA endonuclease as a direct functional target of Mn^2+^. These results support a model in which Mn^2+^ interferes with polymerase-driven RNA synthesis through direct modulation of PA enzymatic function, thereby accounting for the observed pan-subtype antiviral activity. The EDTA chelation–rescue experiment further demonstrates that Mn^2+^ is essential for PA enzymatic activity yet becomes inhibitory at elevated concentrations, supporting a dual regulatory model in which the local Mn^2+^ concentration dictates the functional outcome at the PA active site **(Fig. S3)**. Our experiments with BXA further revealed that its inhibitory effect on PA was strictly dependent on the presence of Mn^2+^: in the absence of Mn^2+^, BXA showed almost no inhibition of PA endonuclease activity, and maximal inhibition was observed only when BXA was pre-incubated with PA prior to the addition of Mn^2+^ **(Fig. S4)**. This finding is consistent with the established mechanism of BXA-mediated PA inhibition through chelation of the binuclear metal center, and provides indirect evidence that Mn^2+^ binds to the PA endonuclease active site.

In vivo, the antiviral efficacy of MnCl_2_ displayed a pronounced dependence on the route of administration. Intranasal delivery conferred significant protection against lethal H1N1 challenge. In preliminary experiments, systemic administration via intraperitoneal injection showed no appreciable protective effect. We speculate that intranasal administration achieves effective local Mn^2+^ concentrations in the respiratory tract, suppressing post-entry viral replication within infected cells and thereby limiting dissemination to the lower airway.

The post-entry mechanism of action established in our cell-based assays also prompted us to explore whether MnCl_2_ could exert therapeutic effects when administered after infection had already been initiated—a scenario more relevant to clinical practice. As a preliminary test of this hypothesis, we delivered MnCl_2_ intranasally at 12 h post-infection in a small cohort (n = 3 per group). Under this delayed-dosing regimen, a trend toward attenuated body weight loss was observed compared with the virus control group **(Fig. S5)**. Although these data are hypothesis-generating rather than confirmatory due to the limited sample size, they are consistent with the post-entry mechanism and provide a rationale for future studies with optimized delayed-dosing protocols and larger cohorts to determine whether Mn^2+^ possesses a genuine therapeutic window.

Several limitations of this study should be acknowledged. First, the precise molecular basis of PA inhibition by Mn^2+^ remains to be resolved structurally. Second, we cannot exclude contributions from other polymerase subunits or host cofactors to the observed antiviral activity. Third, the delayed-treatment observation described above was based on a limited sample size; therefore, the observed trend awaits confirmation in larger cohorts.

In summary, our study identifies Mn^2+^ as a modulator of influenza virus replication that acts, at least in part, by directly inhibiting the PA endonuclease of the viral RNA polymerase. The broad-spectrum activity across H1N1 and H3N2 subtypes, together with the metal ion selectivity and the conserved nature of the PA active site, suggests that this metal-based antiviral mechanism may be applicable to diverse influenza A viruses. These findings expand our understanding of metal ion–virus interactions and provide a conceptual basis for exploring metal-dependent mechanisms as antiviral strategies. Furthermore, the emerging field of precision nutrition, which emphasizes personalized dietary interventions based on individual variations, may offer a broader framework for considering how host manganese status influences susceptibility to viral infections. This represents a promising, albeit underexplored, direction for future research at the interface of nutrition and antiviral immunity.

## Acknowledgement

Author contributions:

Yueqi Wu: Investigation, Methodology, Visualization, Writing – original draft, Writing – review & editing.

Shan Xu: Conceptualization, Supervision, Writing – review & editing.

Yigang Tong: Conceptualization, Supervision.

## Competing interests

All authors declare that they have no competing interests.

Data and materials availability

All data are available in the paper or in the supplementary materials.

## Supplementary Materials

Materials and Methods

Fig. S1 to S5 & Table S1

## Materials and Methods

### 1. Cells and Virus

Madin-Darby canine kidney cells (MDCK) and A549 cells (National Infrastructure of Cell line Resources BMCR) were cultured in Dulbecco’s Modified Eagle’s Medium (DMEM/HIGH GLUCOSE, HyClone), supplemented with 10% fetal bovine serum (FBS, Procell Life Science & Technology Co., Ltd (Wuhan, China)) and 1% penicillin-streptomycin (PS, HyClone) at 37°C in 5% CO_2_. H1N1 virus (A/PR/8/34, ATCC VR-1469) and H3N2 virus (A/Aichi/2/68, ATCC VR-1680) were propagated in MDCK cells. Virus was diluted in PBS containing 0.2% BSA and inoculated onto MDCK monolayers. After adsorption at 37°C for 2 h, the inoculum was removed and replaced with DMEM containing 2 μg/mL TPCK-trypsin, 1% penicillin-streptomycin, and 0.2% BSA. Cells were cultured at 37°C in 5% CO_2_ for 48 h. The supernatant was collected, centrifuged at 3,000 rpm for 10 min at 4°C, aliquoted, snap-frozen in liquid nitrogen, and stored at −80°C.

### 2. Reagents and Treatment

Manganese chloride (MnCl_2_, analytical grade, MedChemExpress) was dissolved in DMSO to prepare a 100 mM stock solution for cell-based assays and a 360 mM stock solution for mouse experiments. Baloxavir marboxil (TargetMol) was dissolved in DMSO to prepare a 10 mM stock solution. Calcium chloride anhydrous (CaCl_2_, RHAWN), copper sulfate (CuSO_4_, RHAWN), zinc chloride (ZnCl_2_, RHAWN), and Magnesium sulfate heptahydrate (MgSO_4_·7H_2_O, Solarbio) were dissolved in deionized water to prepare 100 mM stock solutions and were freshly prepared before each experiment. All stock solutions were diluted in culture medium to the indicated working concentrations before use. The final DMSO concentration in all cell-based assays was kept below 0.1% (v/v).

### 3. Antiviral Activity and Cytotoxicity Assay

MDCK cells were seeded in 96-well plates and cultured overnight. Cells were infected with H1N1 or H3N2 (100 TCID_50_) in the presence of the indicated concentrations of MnCl_2_. After adsorption at 37°C for 1 h, the inoculum was removed, and cells were washed once with PBS. DMEM containing 2 μg/mL TPCK-trypsin, 1% penicillin-streptomycin, 0.2% BSA, and the same concentrations of MnCl_2_ was added, and cells were cultured at 37°C for 24 h. The supernatant was collected, and total viral RNA was extracted using the Super FastPure Cell RNA Isolation Kit (RC102-01, Vazyme). Reverse transcription was performed using HiScript II Q RT SuperMix for qPCR (+gDNA wiper) (R223-01, Vazyme). qPCR was carried out using Taq Pro Universal SYBR qPCR Master Mix (Q712-02, Vazyme) on a QuantStudio 1 Real-Time PCR System. The primer sequences were: IVA-F, 5′-GACCAATCCTGTCACCTCTGAC-3′; IVA-R, 5′-GGGCATTTTGGACAAAGCGTCTACG-3′. The viral inhibition rate was calculated as: Inhibition (%) = (viral RNA copies in virus control group − viral RNA copies in compound-treated group) / viral RNA copies in virus control group × 100%. EC_50_ values were calculated using GraphPad Prism 9. Cytotoxicity was assessed by CCK-8 assay (BS350C, Biosharp) according to the manufacturer’s instructions. CC_50_ values were calculated using GraphPad Prism 9.

### 4. Western blot analysis

Cells or lung tissues were lysed in RIPA buffer containing protease and phosphatase inhibitors. Protein concentrations were determined by BCA assay. Equal amounts of protein were separated by SDS-PAGE and transferred onto PVDF membranes. Membranes were blocked with 5% non-fat milk in TBST for 1 h at room temperature and incubated with primary antibodies against PA (GTX636828, GeneTex), NP (GTX125989, GeneTex), or β-actin (ABL1010, Abbkine) overnight at 4°C. After washing with TBST, membranes were incubated with HRP-conjugated goat anti-rabbit IgG (A21020, Abbkine) for 1 h at room temperature. Bands were visualized by ECL and quantified using ImageJ.

### 5. Immunofluorescence staining

MDCK or A549 cells were seeded on glass coverslips in 24-well plates and cultured overnight. Cells were infected with H1N1 (100 TCID_50_) and treated with MnCl_2_ as indicated. At 24 h post-infection, cells were fixed with 4% paraformaldehyde for 15 min, permeabilized with 0.5% Triton X-100 in PBS for 10 min, and blocked with 5% BSA in PBS for 1 h at room temperature. Cells were incubated with primary antibodies against PA (GTX636828, GeneTex) or NP (GTX125989, GeneTex) overnight at 4°C, followed by DyLight 488-conjugated goat anti-rabbit IgG (A23220, Abbkine) for 1 h at 37°C in the dark. Nuclei were counterstained with DAPI using Antifade Mounting Medium (S2110, Solarbio). Images were acquired using a fluorescence microscope at ×200 magnification.

### 6. Time-of-Addition Assay

To determine the stage of the influenza virus life cycle targeted by MnCl_2_, a time-of-addition (TOA) assay was performed. MDCK cells were seeded to appropriate confluence and subjected to virucidal, attachment/entry, post-entry, whole-course treatment, virus control, and cell control conditions. For the virucidal group, MnCl_2_ was preincubated with H1N1 influenza virus at 37 °C for 1 h before being added to cells for an additional 1 h, after which the inoculum was removed and replaced with fresh growth medium without MnCl_2_. For the attachment/entry group, MnCl_2_ and H1N1 virus were separately preincubated at 37 °C for 1 h and then simultaneously added to cells for 1 h, followed by replacement with MnCl_2_-free growth medium. For the post-entry group, H1N1 virus was preincubated at 37 °C for 1 h and added to cells for 1 h, after which the inoculum was removed and replaced with growth medium containing MnCl_2_. For the whole-course treatment group, MnCl_2_ was preincubated with H1N1 virus at 37 °C for 1 h, added to cells for 1 h, and subsequently replaced with growth medium containing the same concentration of MnCl_2_. Virus control cells were infected with H1N1 virus and cultured in MnCl_2_-free medium, whereas cell control cells were treated throughout with virus-free and MnCl_2_-free medium. Cells and supernatants were collected at the indicated time points post infection for viral titration, protein quantification, and other related analyses.

### 7. Strand-specific quantification of viral mRNA, vRNA, and cRNA

Total RNA was extracted from infected MDCK cells at 2, 4, and 12 h post-infection using the same RNA extraction kit as described above. Strand-specific reverse transcription was performed using the TransScript One-Step gDNA Removal and cDNA Synthesis SuperMix (AT311, TransGen Biotech) with tagged strand-specific primers targeting viral mRNA, cRNA, and vRNA. qPCR was carried out as described above with the corresponding strand-specific primer sets. Relative RNA levels were normalized to GAPDH and calculated using the 2⁻ΔΔCt method. All primer sequences are listed in **Table S1**.

### 8. Mouse Infection Model

Five to six weeks old SPF female BALB/c mice (Beijing Vital River Laboratory Animal Technology Co., Ltd.) were anesthetized and intranasally infected with LD_50_ (lethal dose 50%) of H1N1 virus. Groups included a healthy control, virus control, Baloxavir marboxil-treated and MnCl_2_-treated. For treatment groups, MnCl_2_ (5 mg/kg) or baloxavir marboxil (2 mg/kg) was administered once at the time of viral inoculation. MnCl_2_ was delivered intranasally, whereas baloxavir marboxil was administered by oral gavage. Body weight and survival were monitored daily for 11 days post infection. On day 5 post infection, subsets of mice were euthanized, and lung tissues were collected for viral titration and RT–qPCR analysis. RNA was extracted from mouse lung tissues using TRIzol reagent (Invitrogen, Carlsbad, CA, USA) according to the manufacturer’s instructions. RT-qPCR was carried out as described above. For histopathological analysis, lung tissues were fixed in 10% formalin for 72 h, dehydrated through a graded ethanol series, and embedded in paraffin. Sections (5 μm) were stained with hematoxylin and eosin (H&E). Immunohistochemistry was performed on paraffin-embedded lung sections using an anti-PA antibody (GTX636828, GeneTex) according to standard procedures.

### 9. Expression and Purification of PA Protein

The recombinant plasmid encoding full-length wild-type PA was constructed and maintained in our laboratory, and its sequence was confirmed by first-generation sequencing. The recombinant plasmid was transformed into *E. coli* BL21(DE3) competent cells. Transformants were grown in Luria-Bertani (LB) medium supplemented with 50 µg/mL kanamycin at 37°C with shaking at 200 rpm until the optical density at 600 nm (OD600) reached 0.8. Protein expression was induced by the addition of isopropyl-β-D-thiogalactopyranoside (IPTG) to a final concentration of 0.4 mM, and the culture was incubated at 16°C with shaking at 150 rpm for 24 h. Cells were harvested by centrifugation at 6,000 rpm for 10 min and resuspended in lysis buffer consisting of 20 mM Tris-HCl (pH 8.0), 500 mM NaCl, 0.5% Triton X-100, 20 mg/L RNase A, 20 mg/L DNase I, 10% (v/v) glycerol, 1 mM DTT, and 1× protease inhibitors. Cell disruption was achieved using a cell crusher, and insoluble debris was removed by centrifugation at 18,000 rpm for 45 min at 4°C. The cleared supernatant was incubated with Ni-NTA agarose at 4°C for 4 h to allow binding of the His-tagged protein. The resin was sequentially washed with wash buffer (20 mM Tris-HCl pH 8.0, 500 mM NaCl, 0.4 mM DTT, 30 mM imidazole) and high-salt wash buffer containing 60 mM imidazole. Target proteins were eluted with elution buffer (20 mM Tris-HCl pH 8.0, 500 mM NaCl, 500 mM imidazole, 1 mM DTT, 10% glycerol). The eluate was concentrated using 30 kDa centrifugal filters and dialyzed against dialysis buffer (20 mM Tris-HCl pH 8.0, 100 mM NaCl, 10% glycerol, 1 mM DTT). Purity of the purified protein was assessed by SDS-PAGE followed by Coomassie brilliant blue staining. Aliquots of the purified protein were flash-frozen in liquid nitrogen and stored at −80°C until further use.

### 10. PA endonuclease activity assay

The endonuclease activity of recombinant PA protein was assessed using two complementary detection methods: denaturing polyacrylamide gel electrophoresis (PAGE) and fluorescence resonance energy transfer (FRET). The RNA substrate (5′-6-FAM-CUCCUCAUUUUUCCCUAGUU-BHQ1-3′) was used for both assays.

For the gel-based assay, the reaction mixture (10 μL total volume) contained 1 μL of 20-fold diluted purified PA protein (original concentration 0.503 mg/mL), 2 µM fluorophore-labeled RNA substrate, and reaction buffer (20 mM Tris-HCl, 100 mM NaCl, pH 8.0). After incubation at 37°C for 10 min, the reaction was terminated by adding EGTA (final concentration 20 mM). For inhibition assays, PA protein was pre-incubated with the test compound at 37°C for 15 min prior to the addition of the substrate. Cleavage products were resolved on a 25% urea-denaturing polyacrylamide gel (Solarbio Nucleic Acid Denaturing PAGE Preparation Kit, P1330) at 270 V for 60 min and visualized using a fluorescence imaging system.

For the fluorescence-based assay, the reaction mixture (50 μL total volume) contained 15-fold diluted purified PA protein and 750 nM fluorophore-labeled RNA substrate in reaction buffer (20 mM Tris-HCl, 100 mM NaCl, pH 8.0). PA protein was pre-incubated with the indicated concentrations of MnCl_2_ at 37°C for 15 min, and the reaction was initiated by adding the substrate. Fluorescence intensity was monitored continuously at room temperature using a BioTek Synergy H1 microplate reader with excitation at 465 nm and emission at 520 nm. The increase in fluorescence reflected endonucleolytic cleavage of the substrate. Reaction rates were calculated from the fluorescence signals at the 10-min time point, and inhibition rates were determined relative to the MnCl_2_-free PA control. IC_50_ values were calculated by nonlinear regression using GraphPad Prism 9.

### 11. EDTA chelation–rescue assay

Purified PA protein was pre-treated with EDTA (final concentration 5 mM) at 37°C for 10 min to chelate active-site metal ions. The indicated concentrations of MnCl_2_ (100 μM, 500 μM, or 1 mM) were then added, and the mixture was incubated at 37°C for an additional 10 min. The reaction was initiated by adding a 5′-FAM / 3′-BHQ1 dual-labeled RNA substrate (final concentration 750 nM). Untreated PA and EDTA-treated PA without Mn^2+^ rescue served as controls. Fluorescence intensity was monitored continuously for 45 min at room temperature using a BioTek Synergy H1 microplate reader with excitation at 465 nm and emission at 520 nm. The reaction mixture (50 μL total volume) contained 15-fold diluted purified PA protein in reaction buffer (20 mM Tris-HCl, 100 mM NaCl, pH 8.0).

### 12. Ethics statement

All animal procedures were conducted under protocols approved by the Animal Care and Use Committee of Ocean University of China.

### 13. Statistical analysis

All data are representative of at least three independent experiments and expressed as the mean ±SD. The data was analyzed with the GraphPad Prism 9 software to evaluate the statistical significance by two-tailed unpaired Student’s t-test. A statistically significant difference was accessed when P < 0.05. P < 0.05 was marked *. P < 0.01 was marked **, P < 0.001 was marked *** and P < 0.0001 was marked ****.

---

**Fig. S1.**
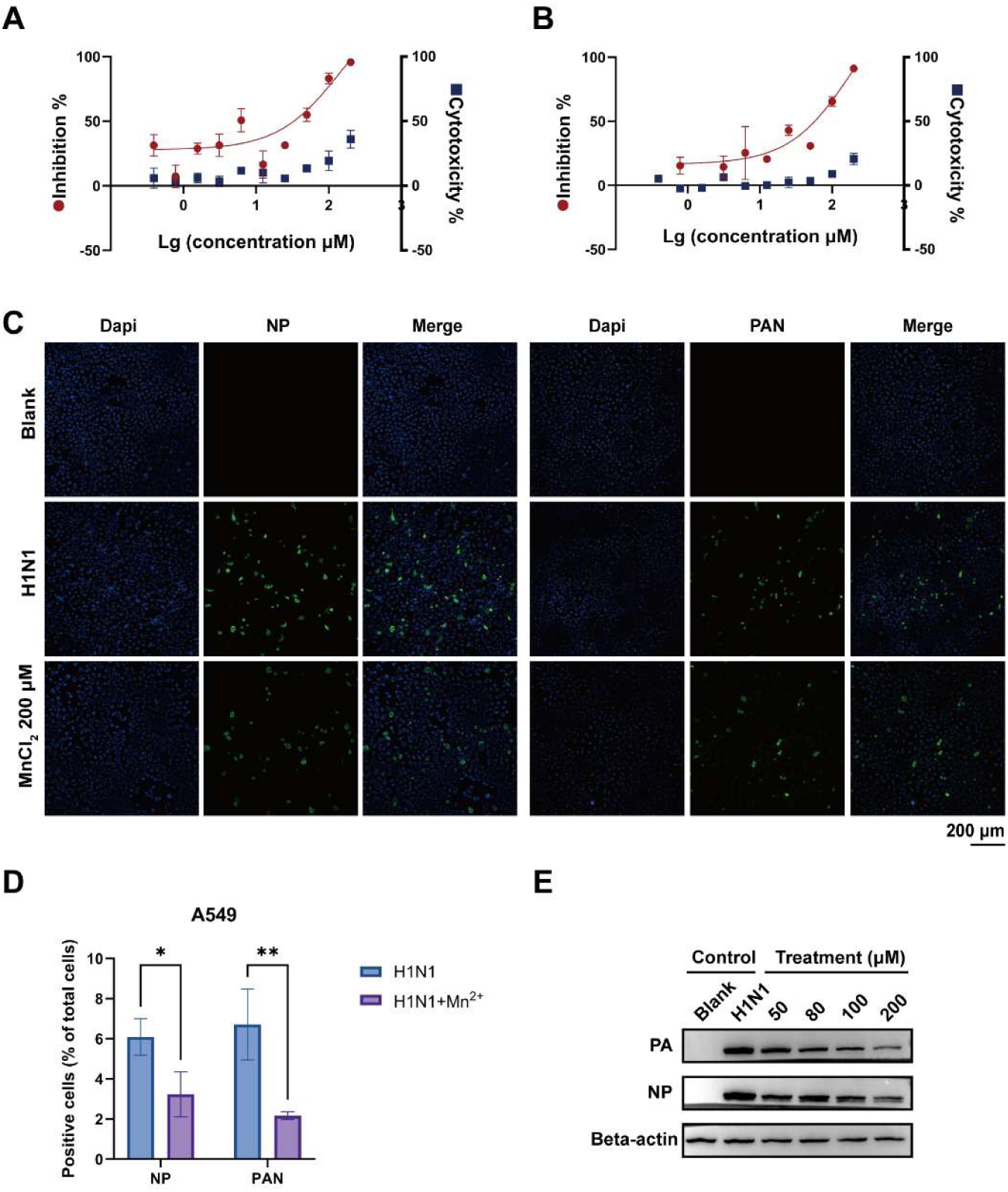
**Mn**^2+^ **inhibits H1N1 and H3N2 replication in A549 cells. (A)** Antiviral activity (red) and cytotoxicity (blue) of Mn^2+^ against H1N1 in A549 cells. **(B)** Antiviral activity (red) and cytotoxicity (blue) of Mn^2+^ against H3N2 in A549 cells. **(C)** Immunofluorescence staining of viral NP (green) and PA (green) in H1N1-infected A549 cells with or without Mn^2+^ (200 μM) treatment. **(D)** Quantification of NP- and PA-positive cells from (C). **(E)** Western blot analysis of NP and PA protein levels in H1N1-infected A549 cells treated with Mn^2+^. β-actin served as loading control. **Data information:** In (A, B, D), data are presented as mean ± SD (n = 3). *P < 0.05, **P < 0.01 versus virus-infected control (Student’s t-test).

**Fig. S2.**
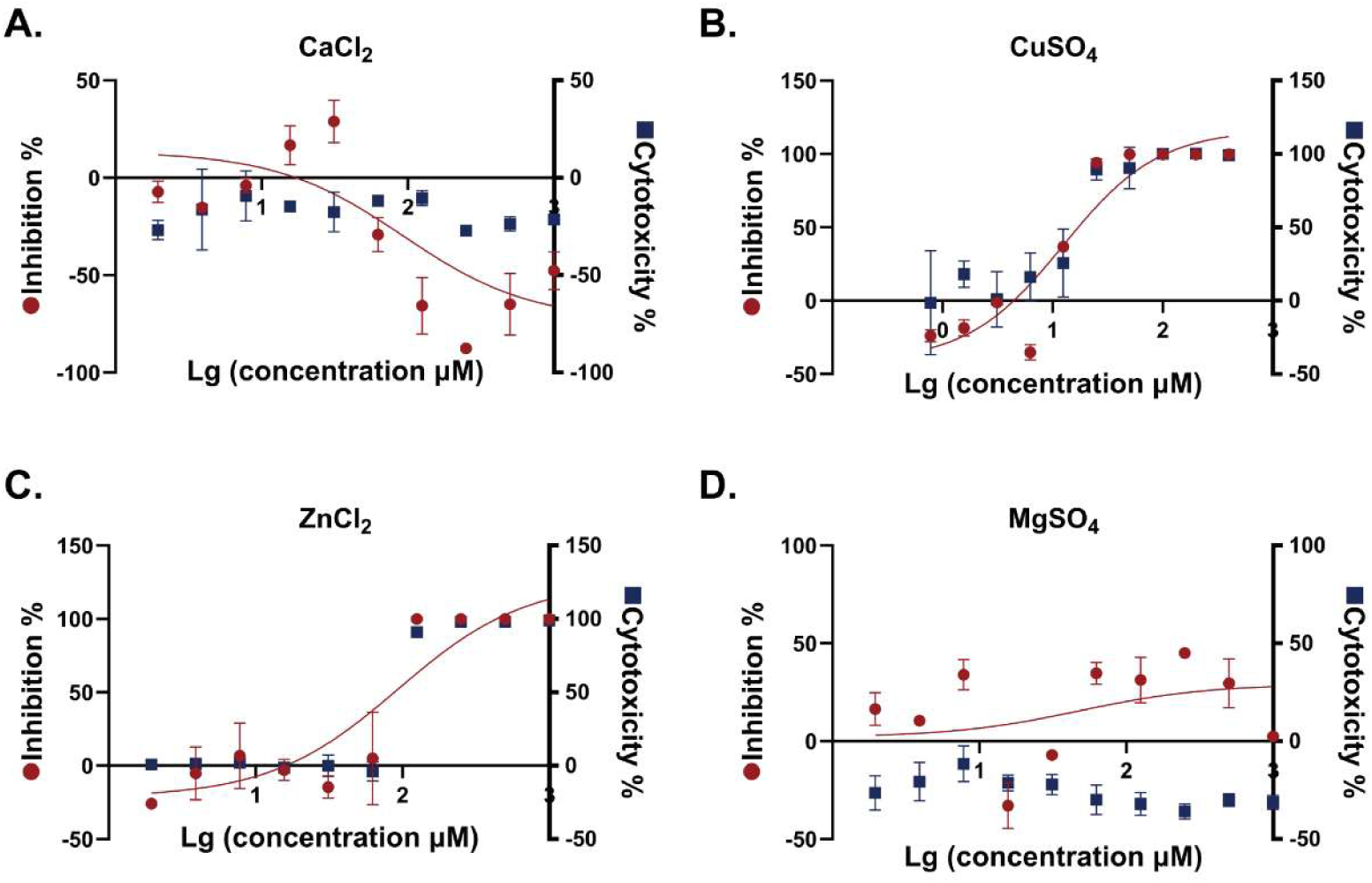
Effect of other divalent metal ions on H1N1 replication in MDCK cells. (A-D) Antiviral activity (red) and cytotoxicity (blue) of Ca^2+^, Cu^2+^, Zn^2+^, or Mg^2+^. **Data information:** Data are presented as mean ± SD (n = 3).

**Fig. S3.**
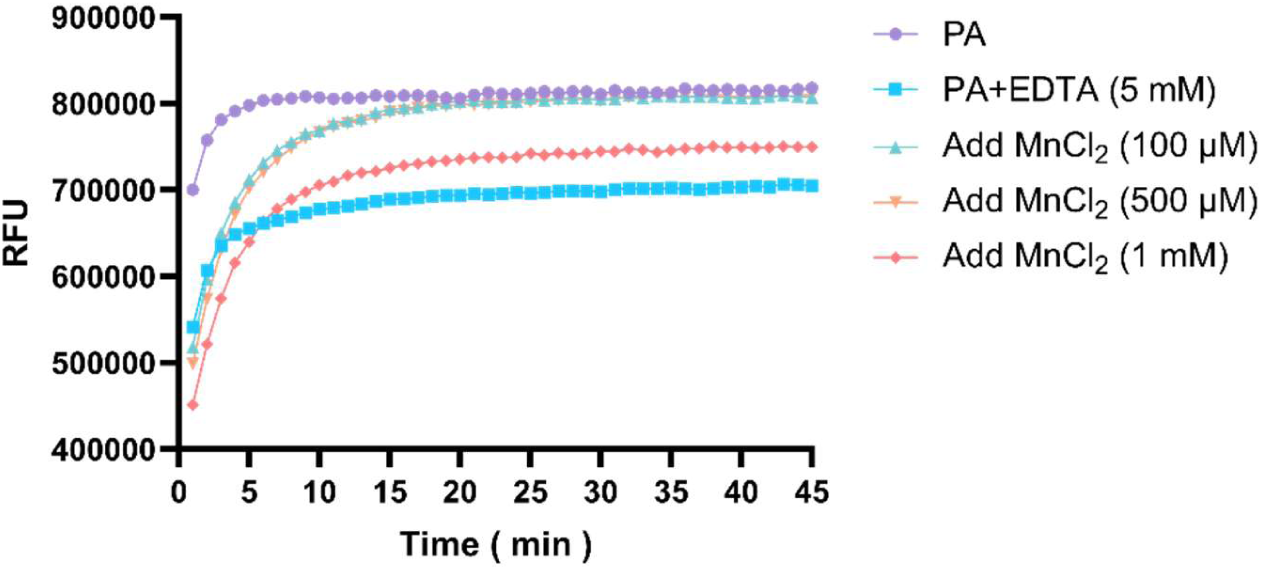
EDTA chelation–rescue assay reveals dual regulation of PA endonuclease activity by Mn^2+^. PA protein was pre-treated with EDTA and then supplemented with the indicated concentrations of MnCl_2_. Untreated PA and EDTA-treated PA without Mn^2+^ rescue served as controls. Fluorescence intensity was monitored continuously for 45 min.

**Fig. S4.**
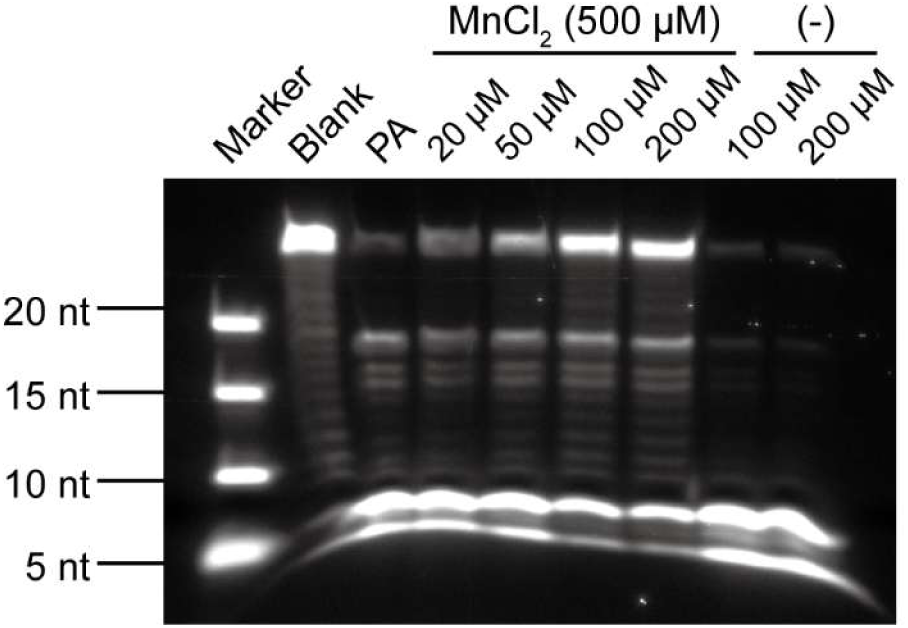
BXA inhibits PA endonuclease in a Mn^2+^-dependent manner. Purified PA protein was incubated with BXA in the absence or presence of Mn^2+^. Untreated PA and BXA-only controls were included. Reaction products were resolved by denaturing PAGE.

**Fig. S5.**
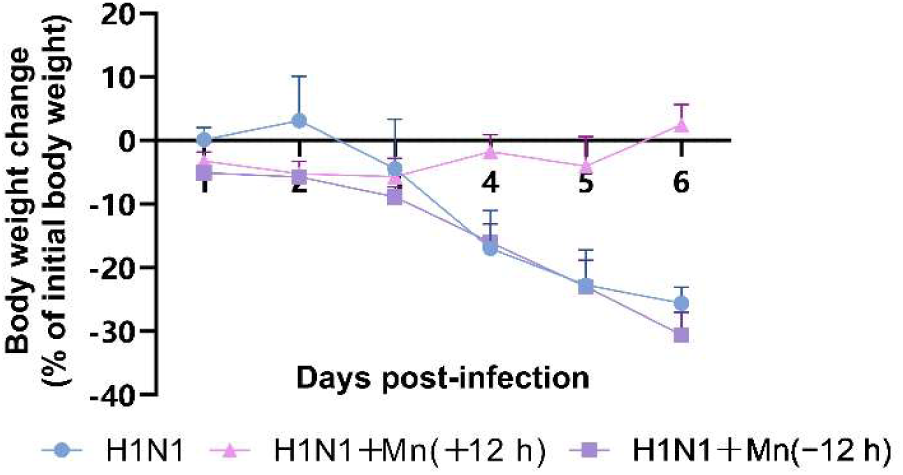
Effect of delayed MnCl_2_ treatment on body weight change in H1N1-infected mice (preliminary). BALB/c mice (n = 3 per group) received a single intranasal dose of MnCl_2_ (5 mg/kg) or PBS (virus control) at 12 h post-infection. Body weight was normalized to the initial body weight on day 0. **Data information:** Data are presented as mean ± SD (n = 3). No statistical test was applied to this preliminary experiment.

**Table S1.**
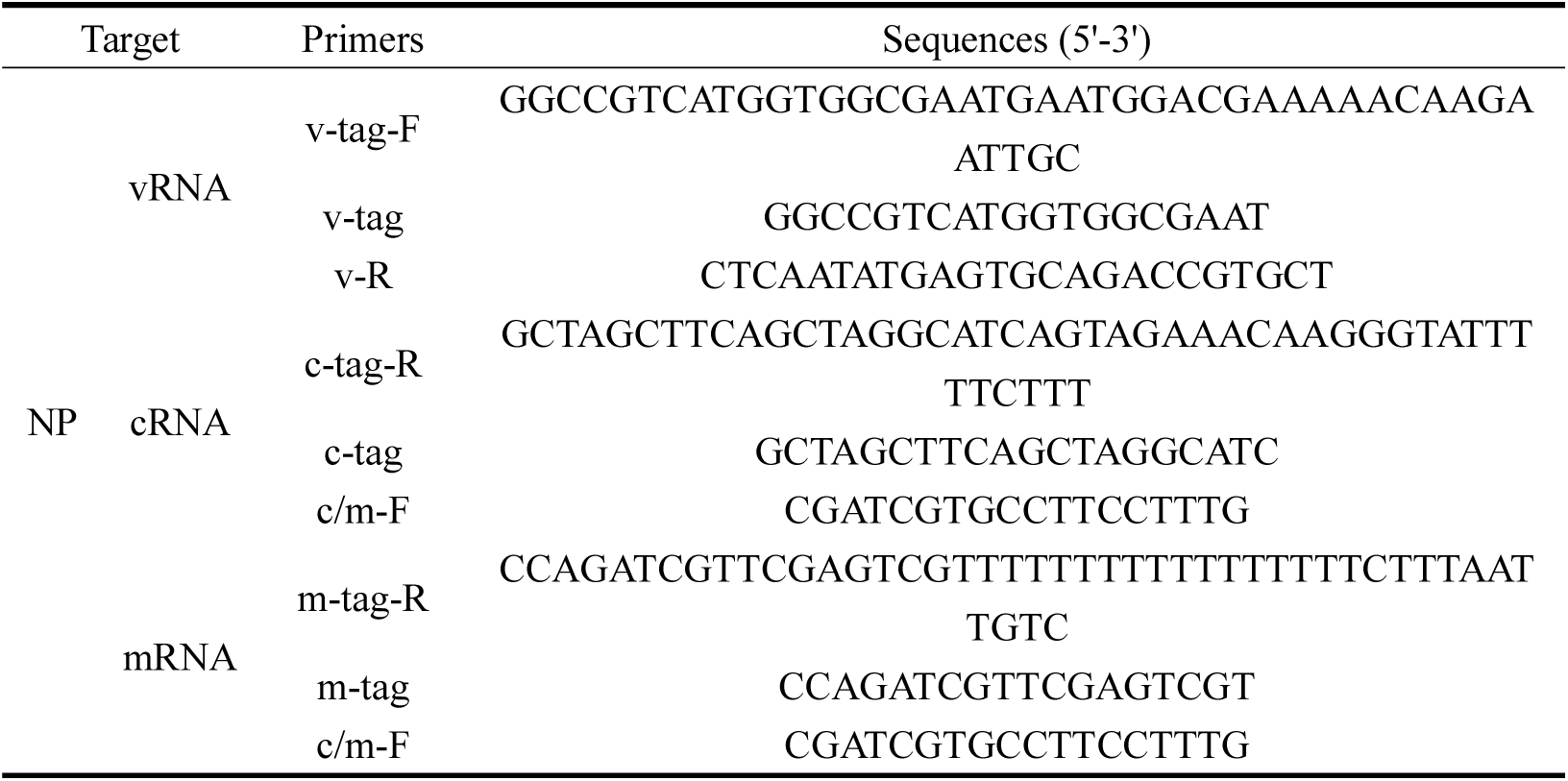
Primers used for strand-specific RT-qPCR.

